# Characterizing the temporal dynamics of gene expression in single cells with sci-fate

**DOI:** 10.1101/666081

**Authors:** Junyue Cao, Wei Zhou, Frank Steemers, Cole Trapnell, Jay Shendure

## Abstract

Gene expression programs are dynamic, *e.g.* the cell cycle, response to stimuli, normal differentiation and development, etc. However, nearly all techniques for profiling gene expression in single cells fail to directly capture the dynamics of transcriptional programs, which limits the scope of biology that can be effectively investigated. Towards addressing this, we developed *sci-fate*, a new technique that combines S4U labeling of newly synthesized mRNA with single cell combinatorial indexing (sci-), in order to concurrently profile the whole and newly synthesized transcriptome in each of many single cells. As a proof-of-concept, we applied sci-fate to a model system of cortisol response and characterized expression dynamics in over 6,000 single cells. From these data, we quantify the dynamics of the cell cycle and glucocorticoid receptor activation, while also exploring their intersection. We furthermore use these data to develop a framework for inferring the distribution of cell state transitions. We anticipate sci-fate will be broadly applicable to quantitatively characterize transcriptional dynamics in diverse systems.

## Main

During organismal development, as well as during myriad physiological and pathophysiological processes, individual cells traverse a manifold of molecularly and functionally distinct states. The accurate characterization of these trajectories is key to advancing our understanding of each such process, for identifying the causal factors that drive them, and for rationally designing effective perturbations of them. However, although experimental methods for profiling various aspects of single cell biology have recently proliferated, nearly all such methods deliver only a static snapshot of each cell, *e.g.* of gene expression at the moment of fixation.

To recover temporal dynamics, several groups have developed computational methods that place individual cells along a continuous trajectory based on single cell RNA-seq data, *i.e.* the concept of pseudotime^1–6^. However, such methods are inherently limited in several important ways, including that they are inferring rather than directly measuring dynamics, that they are dependent on sufficient representation across the trajectory, and that they may fail to capture the detailed dynamics of individual cells (*e.g.* directionality, multiple superimposed potentials, etc.)^7^. Although time-lapse microscopy is a distinct technology that overcomes some of these limitations, it is limited in throughput and scope (*e.g.* enabling visualization of a few marker genes in a few cells), and as such may be insufficient to decipher the complexity of many biological systems.

Here we describe a novel technique, sci-fate, to measure the dynamics of gene expression in single cells at the level of the whole transcriptome. In brief, we integrated protocols for labeling newly synthesized mRNA with 4-thiouridine (S4U)^8,9^ with single cell combinatorial indexing RNA-seq (sci-RNA-seq^10^). As a proof-of-concept, we applied sci-fate to a model system of cortisol response, and characterized expression dynamics in over 6,000 single cells. From these data, we quantify the dynamics of the transcription factor modules that underpin the cell cycle, glucocorticoid receptor activation, and other processes. We furthermore use these data to develop a framework for inferring the distribution of cell state transitions. The experimental and computational methods described here may be broadly applicable to quantitatively characterize transcriptional dynamics in diverse systems.

### Overview of sci-fate

Briefly, sci-fate relies on the following steps (Fig. 1a): (i) Cells are incubated with 4-thiouridine (S4U), a thymidine analog, to label newly synthesized RNA^11–17^. (ii) Cells are harvested, fixed with 4% paraformaldehyde, and then subjected to a thiol(SH)-linked alkylation reaction which covalently attaches a carboxyamidomethyl group to S4U by nucleophilic substitution ^8^. (iii) Cells are distributed by dilution to 4 x 96 well plates. The first sci-RNA-seq molecular index is introduced via *in situ* reverse transcription (RT) with a poly(T) primer bearing both a well-specific barcode and a degenerate unique molecular identifier (UMI). During first strand cDNA synthesis, modified S4U templates guanine rather than adenine incorporations. (iv) Cells from all wells are pooled and then redistributed by fluorescence-activated cell sorting (FACS) to multiple 96-well plates. Cells are gated on DAPI (4’,6-diamidino-2-phenylindole) to distinguish singlets from doublets. (v) Double-stranded cDNA is generated by RNA degradation and second-strand synthesis. After Tn5 transposition, cDNA is PCR amplified via primers recognizing the Tn5 adaptor on the 5’ end and the RT primer on the 3’ end. These primers also bear a well-specific barcode that introduces the second sci-RNA-seq molecular index. (vi) PCR amplicons are subjected to massively parallel DNA sequencing. As with other sci-methods^18–25^, most cells pass through a unique combination of wells, such that their contents are marked by a unique combination of barcodes that can be used to group reads derived from the same cell. (vii) The subset of each cell’s transcriptome corresponding to newly synthesized transcripts is distinguished by T→ C conversions in reads mapping to mRNAs (**Methods**).

**Fig. 1.**
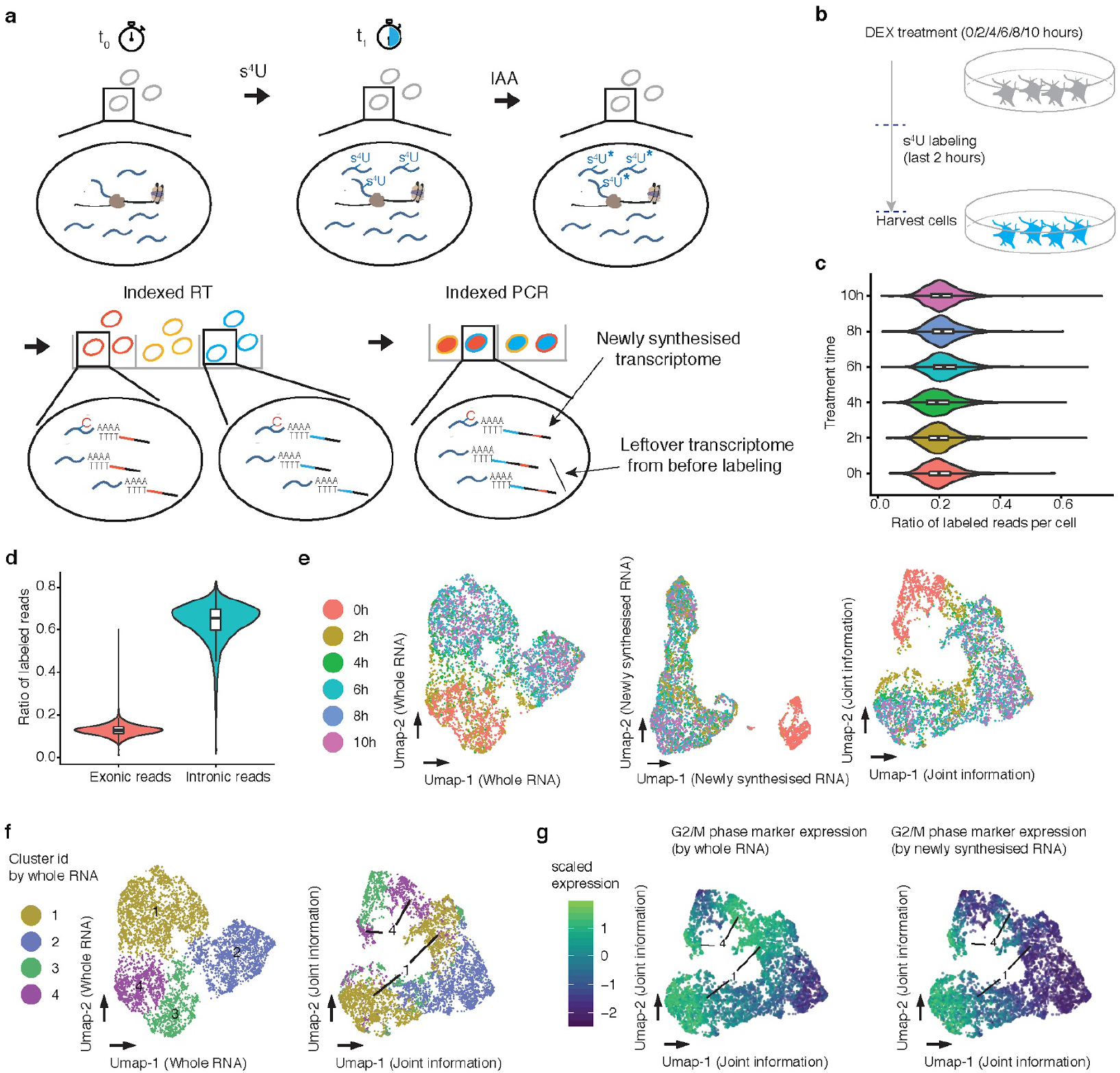
Sci-fate enables joint profiling of whole and newly synthesized transcriptomes. (**a**) The sci-fate workflow. Key steps are outlined in text. (**b**) Experimental scheme. A549 cells were treated with dexamethasone for varying amounts of time ranging from 0 to 10 hrs. Cells from all treatment conditions were labeled with S4U two hours before harvest for sci-fate. (**c**) Violin plot showing the fraction of S4U labeled reads per cell for each of the six treatment conditions. For all violin plots in this figure: thick horizontal lines, medians; upper and lower box edges, first and third quartiles, respectively; whiskers, 1.5 times the interquartile range; circles, outliers. (**d**) Violin plot showing the fraction of S4U labeled reads per cell, split out by the subsets that map to exons vs. introns. (**e**) UMAP visualization of A549 cells based on their whole transcriptomes (left), newly synthesized transcriptomes (middle) or with joint analysis, *i.e.* combining the top PCAs from each (right). (**f**) Same as left and right of panel E, respectively, but colored by cluster id from UMAP based on whole transcriptomes. (**g**) Same as right of panel E, but colored by normalized expression of G2/M marker genes by their overall expression levels (left) or their levels of newly synthesized transcripts (right). UMI counts for these genes are scaled by library size, log-transformed, aggregated and then mapped to Z-scores.

For quality control, we first tested sci-fate with a mixture of HEK293T (human) and NIH/3T3 (mouse) cells under four conditions: with vs. without S4U labeling (200 nM, 6 hrs), and with vs. without the thiol(SH)-linked alkylation reaction. Under all four conditions, transcriptomes from human/mouse cells were overwhelmingly species-coherent (>99% purity for both human and mouse cells, 2.7% collisions; Extended Data Fig. 1ab) with similar mRNA recovery rates (overall median 21,342 UMIs per cell; Extended Data Fig. 1c). However, only with S4U labeling and the thiol(SH)-linked alkylation reaction did we observe a substantial proportion of reads bearing one or more T → C conversions, *i.e.* newly synthesized transcripts (46% for human and 31% for mouse cells, as compared with 0.8% for both species in untreated cells; Extended Data Fig. 1d). The aggregated transcriptomes of cells derived from sci-fate and conventional sci-RNA-seq on the same cell types were highly correlated (Spearman’s correlation r = 0.99; Extended Data Fig. 1ef), suggesting that the short term labeling and conversion process do not substantially bias transcript counts.

### Joint profiling of the total and newly synthesized transcriptome in cortisol response

Cortisol influences the activity of almost every cell in the body, regulating genes involved in diverse processes including development, metabolism and immune response^26^. To investigate the dynamics of cortisol response, we applied sci-fate to an *in vitro* model wherein dexamethasone (DEX), a synthetic mimic of cortisol, activates glucocorticoid receptor (GR), which binds to thousands of locations across the genome and significantly alters cell state within a rapid timeframe^27–30^. Specifically, we treated lung adenocarcinoma-derived A549 cells for 0, 2, 4, 6, 8 or 10 hrs with 100 nM DEX. In each condition, cells were incubated with S4U (200 nM) for the two hours immediately preceding harvest. We then performed 384 x 192 sci-fate (Fig. 1b). Each of the six conditions was represented by 64 wells during the first round of indexing, such that all samples could be processed in a single sci-RNA-seq experiment to minimize batch effects.

After filtering out low quality cells, potential doublets and a small subgroup of differentiated cells (**Methods**), we obtained single cell profiles for 6,680 cells (median of 26,176 UMIs corresponding to mRNAs detected per cell). A median of 20% of mRNA UMIs were labeled per cell (Fig. 1c; Extended Data Fig. 2a-c). The proportion of newly synthesized mRNAs was markedly higher in reads mapping to intronic (65%) vs. exonic (13%) regions (p-value < 2.2e-16, Wilcoxon signed rank test; Fig. 1d), consistent with the expectation that the intronic reads are more likely to have been recently synthesized.

In exploring these data, we first asked whether the newly synthesized vs. whole transcriptome data convey identical or distinct information with respect to cell state. For each condition, we generated pseudobulk transcriptomes for either the newly synthesized or whole transcriptomes (*i.e.* aggregating across cells), and compared these in a pairwise fashion between conditions (*e.g.* whole transcriptome at 0 hrs vs. 4 hrs; newly synthesized transcriptome at 2 hrs vs. 6 hrs, etc.) (Extended Data Fig. 2d). The lowest correlations corresponded to the newly synthesized transcriptome with no DEX treatment (0 hrs) vs. the newly synthesized transcriptomes of any DEX treated condition (Extended Data Fig. 2d). Consistent with this, performing dimensionality reduction with Uniform Manifold Approximation and Projection (UMAP)^31^ on whole transcriptomes failed to separate DEX untreated (0 hrs) vs. treated (2+ hrs) cells (Fig. 1e, left). In contrast, applying UMAP to the newly synthesized subset of the single cell transcriptomes readily separated DEX untreated vs. treated cells (Fig. 1e, center). These patterns are likely a consequence of the fact that in DEX treated cells, the newly synthesized transcriptome more faithfully reflects the GR response itself. Illustrative of this, the classic markers for GR response, *FGD4*^27^ and *FKBP5*^32^, exhibit the highest fold induction in comparing the newly synthesized transcriptome at 0 hrs vs. 2 hrs, but the magnitude of their induction is dampened when comparing the whole transcriptome between the same time points (Extended Data Fig. 2ef; **Supplementary Table 1**).

To jointly make use of the information conveyed by the whole and newly synthesized transcriptomes, we combined their top principal components (PCs) for UMAP analysis. This approach separates cells that had experienced no DEX treatment (0 hrs), recent treatment (2 hrs) or extended treatment (4+ hrs) (Fig. 1e, right). Interestingly, with this joint approach, the cells corresponding to two clusters defined by analysis of whole transcriptomes (clusters 1 & 4 in Fig. 1f) each split into two groups (Fig. 1f). By examining the levels of newly synthesized mRNAs corresponding to cell cycle markers^33^, we found that one pair of these new groups corresponds to cells in G2/M phase (high levels of both overall and newly synthesized G2/M markers), and the other to early G0/G1 phase cells (high levels of overall but low levels of newly synthesized G2/M markers) (Fig. 1g; Extended Data Fig. 2gh). These analyses indicate that joint analysis of the newly synthesized and whole single cell transcriptomes can recover cell state information that is not easily obtained from whole transcriptomes alone.

### TF module activity decomposes GR response, cell cycle, and other cellular processes

Multiple dynamic gene regulatory processes are concurrently underway in this *in vitro* GR response system -- minimally, the GR response itself and the cell cycle. We speculated that these might be disentangled, and their intersection probed, by first identifying the transcription factor (TF) modules driving new mRNA synthesis in relation to each such process.

Towards identifying such modules, candidate links between TFs and their regulated genes were identified as follows. For each gene, across the 6,680 cells, we computed correlations between the levels of newly synthesized mRNA for that gene and the overall expression level of each of 859 transcription factors, using LASSO (least absolute shrinkage and selection operator) regression. Out of 1,086 links with TFs characterized by ENCODE^34^, 807 were validated by TF binding sites near the genes’ promoters^34^, a 4.3-fold enrichment relative to background expectation (odds ratio for validation = 2.89 for links identified in LASSO regression vs. 0.67 for background, p-value < 2.2e-16, Fisher’s Exact test). These covariance links were further filtered by ChIP-seq binding^35^ and motif^36^ enrichment analysis (Fig. 2a, **Methods**). In total, we identified 986 links between 29 TFs and 532 genes (Fig. 2ab, **Supplementary Table 2**). As a control analysis, we permuted the cell IDs of the TF expression matrix and repeated the analysis. Under the same thresholds, no links were identified after permutation. Some of the identified TF and gene regulatory relationships are readily validated in a manually curated database of TF networks (TRRUST^37^), such as E2F1 (top enriched TF of E2F1 linked genes = E2F1, adjusted p-value = 8e-7)^38^, NFE2L2 (top enriched TF of NFE2L2 linked genes = NFE2L2, adjusted p-value = 0.003)^38^, and SREBF2 (top enriched TF of SREBF2 linked genes: SREBF2, adjusted p-value = 0.0006)^38^.

**Fig. 2.**
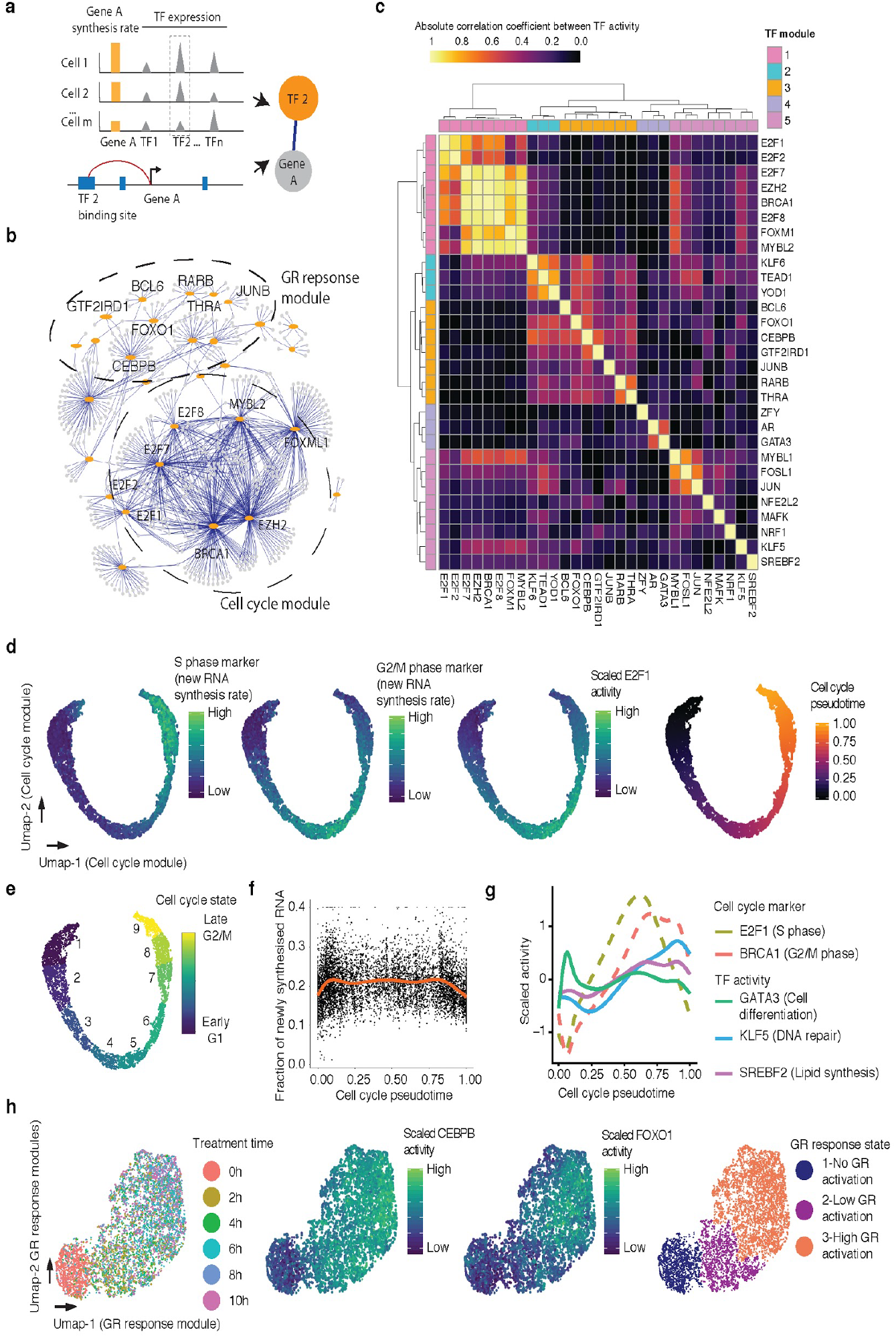
Characterizing TF modules driving concurrent, dynamic gene regulatory processes in populations of single cells. (**a**) Schematic of approach used to identify links between TFs and their regulated genes. (**b**) Identified links (blue) between TFs (orange) and regulated genes (grey). TF modules related to cell cycle progression and GR response are labeled. (**c**) Heatmap showing the absolute Pearson’s correlation coefficient between the activities of pairs of TFs. (**d**) UMAP visualization of A549 cells based on activity of cell cycle-related TF module, colored by levels of newly synthesized mRNA corresponding to, from left to right, S phase markers, G2/M phase markers, and *E2F1* activity. Rightmost panel is colored by pseudotime based on point position on the principal curve estimated by princurve package^54^. (**e**) Same as panel **d**, but colored according to nine cell cycle states defined by unsupervised clustering analysis. In broad terms, cell cycle states 1-3 correspond to G1 phase, 4-6 to S phase, and 7-9 to G2/M phase. (**f**) Scatter plot showing the changes in the fraction of newly synthesized mRNA in each cell along cell cycle progression. The blue line is the smoothed curve estimated by the geom_smooth function^55^. (**g**) Similar to panel F, but showing smoothed activity of selected TF modules as a function of cell cycle pseudotime. (**h**) UMAP visualization of A549 cells based on activity of GR response-related TF module, colored by DEX treatment time (left), *CEBPB* or *FOXO1* activity (middle panels), or cluster id from unsupervised clustering (right). Throughout figure, to calculate TF module activity, newly synthesized UMI counts for genes linked to module-assigned TFs are scaled by library size, log-transformed, aggregated and then mapped to Z-scores.

The 29 TFs with one or more gene links included well-established GR response effectors such as *CEBPB*^*39*^, *FOXO1*^*40*^, and *JUNB*^*41*^ (Fig. 2b; Extended Data Fig. 3ab). This group also included several TFs that have not previously been implicated in GR response, including *YOD1* and *GTF2IRD1*, both of which exhibited greater expression and activity in DEX treated cells (Extended Data Fig. 3cd). The main TFs driving cell cycle progression were also identified, including *E2F1*, *E2F2*, *E2F7*, *BRCA1*, and *MYBL2*^*42*^. Notably, the expression levels of TFs such as *E2F1* were more highly correlated with the levels of newly synthesized target gene mRNA than the overall levels of target gene mRNA (Extended Data Fig. 3e). We also observed regulatory links corresponding to TFs involved in cell differentiation such as *GATA3*^*43*^, mostly expressed in a subset of quiescent cells, as well as TFs involved in oxidative stress response such as *NRF1*^*44*^ and *NFE2L2* (*NRF2*)^45^.

We calculated a measure of each of these 29 TFs’ activities in each cell, based on the normalized aggregation of the levels of newly synthesized mRNA for all of its target genes. We then computed the absolute correlation coefficient between each possible pair of TFs with respect to their activity across the 6,680 cells. Hierarchical clustering of these pairwise correlations resulted in the identification of several major TF modules, *i.e.* sets of TFs that appear to be regulating the same process (Fig. 2c). A first TF module corresponds to all cell cycle-related TFs in the set, *e.g. E2F1* and *FOXM1*^*42*^. A second large module corresponds to GR response-related TFs including *FOXO1*, *CEBPB*, *JUNB* and *RARB*^*39–41*^. The other modules include one corresponding to GR-activated G1/G2/M phase cells (*KLF6*, *TEAD1*, and *YOD1*; Extended Data Fig. 3f), and another corresponding to likely-differentiating GR-activated G1 phase cells *GATA3* and *AR*; Extended Data Fig. 3f)^43,46^. Additional TFs or TF modules appear to capture other processes that are heterogeneous in this population of cells, including *NRF1* and *NFE2L2* for stress response/apoptosis (top enriched pathway of *NFE2L2* linked genes: ferroptosis, adjusted p-value = 1e-5)^38,44,45,47^; *KLF5* for DNA damage repair (top pathway: ATM signaling, adjusted p-value = 0.018)^38,48^; and *SREBF2* for cholesterol homeostasis (top pathway: “SREBF and miR33 in cholesterol and lipid homeostasis”, adjusted p-value = 9e-6)^38,49^.

To assign cell cycle states to individual cells, we first ordered cells by their cell cycle-linked TF module activity. This resulted in a smooth, nearly circular trajectory, in which the levels of newly synthesized mRNA corresponding to known cell cycle markers was dynamic (Fig. 2d)^33^. We observed a gap between late G2/M phase and early G1 phase, consistent with the dramatic cell state change during cell division. By unsupervised clustering of the activities of individual TFs within the cell cycle-linked TF module, we identified 9 cell cycle states spanning the early, middle and late cell cycle phases (Fig. 2e). Early G1 and late G2/M phase cells exhibited decreased synthesis of new RNA relative to other parts of the cell cycle, possibly due to chromosomal condensation during mitosis (Fig. 2f)^50–52^. Other (*i.e.* non-cell-cycle) TF modules exhibited different dynamics in relation to cell cycle progression (Fig. 2g). For example, *GATA3* activity peaks in early G1 phase, potentially reflecting a cell differentiation pathway distinct from cell cycle reentry^43^. In contrast, the modules of *KLF5* and *SREBF2*, associated with DNA repair and lipid homeostasis, respectively, exhibit greater activity from S to G2 phase, possibly related to roles in DNA replication and cell division, respectively^53^.

With similar approaches, the cells can also be ordered into a smooth trajectory based on GR response-linked TF module activity. As expected, this trajectory correlates well with DEX treatment time, as well as the activity of GR response-related TFs (Fig. 2h). By unsupervised clustering of the activities of individual TFs within the GR response-linked TF module, we identified GR response states corresponding to no, low and high levels of activation (Fig. 2h).

We next sought to explore the intersection of the 9 cell cycle states (Fig. 2e) and the three GR response states (Fig. 2h). Each of their 27 possible state combinations were represented by some cells, with the smallest group corresponding to 1.1% of the overall dataset (n = 74 cells, intersection of “early G2/M” cell cycle state and “no GR activation” state; Extended Data Fig. 4ab). Although we observe several TF modules that appear specific to certain intersections of the cell cycle and GR response (*KLF6/TEAD1/YOD1* and *GATA3/AR*, discussed above), several observations support the conclusion that the dynamics of the cell cycle and GR response operate largely independently. First, we observe minimal correlation between the activities of the primary TF modules for cell cycle and GR response across the 6,680 cells (Pearson’s correlation r = 0.004; Fig. 2c). Second, the relative proportions of each of the 27 possible state combinations are readily predicted by proportions of cell cycle and GR response states (Extended Data Fig. 4b).

### Inferring single cell transcriptional dynamics with sci-fate

We next sought to develop a strategy to use sci-fate data to infer the *past* transcriptional state of each cell, *i.e.* at the onset of S4U labeling, which might in turn allow us to relate cells derived from different timepoints. The inference of this past transcriptional state requires knowledge of two parameters -- first, the detection rate of newly synthesized transcripts, and second, the degradation rate of each mRNA species. Below, we discuss how each of these parameters can be estimated directly from the sci-fate data generated for this experiment. A more detailed consideration is provided in the **Methods**.

Under the assumption that mRNA degradation rates are not affected by DEX treatment (this assumption is validated further below), it is relatively straightforward to estimate sci-fate’s detection rate for newly synthesized transcripts. Each sci-fate transcriptome in this dataset consists of two components -- the newly synthesized transcriptome, whose detection rate we are hoping to estimate, and the ‘leftover’ transcriptome, *i.e.* transcripts that were present at the onset of 4SU labeling, minus any degradation over the course of the two hours. Comparing the 0 hr (untreated) and 2 hr DEX treatment groups, we expect that their leftover transcriptomes should be identical, as should sci-fate’s detection rate for newly synthesized transcripts. As such, an equation can be constructed relating the transcriptomes of these treatment groups to one another (**Methods**). For each of 186 genes exhibiting the largest differences in new transcription between the two conditions, we solved this equation to estimate sci-fate’s detection rate. As these estimates were largely consistent across genes (Extended Data Fig. 5ab), we used their median value (82%) as sci-fate’s estimated detection rate for all subsequent analyses.

We next sought to estimate the degradation rate of each mRNA species. As noted above, the bulk transcriptome at each timepoint in our experiment can be decomposed into the newly synthesized transcriptome and the leftover transcriptome. Furthermore, the leftover transcriptome should equal the bulk transcriptome from the timepoint two hours earlier, but corrected for mRNA degradation over that interval. From these assumptions, an equation can be constructed and solved to estimate the mRNA half-life of each gene, which we did independently for each two hour interval of the experiment (**Methods**; **Supplementary Table 3**). As a first quality check, we simply compared these estimated mRNA half-lives between timepoints, and found them to be highly consistent (**Extended Data Fig. 5c**; median Pearson’s *r* = 0.92). As a second quality check, we compared them to orthogonally generated estimates of mRNA half-lives from the literature^9^. Despite the fact that different technologies were used on different cell lines (A549 vs. K562), the estimates of mRNA half-lives were reasonably consistent (Extended Data Fig. 5d; Pearson’s *r* = 0.76).

With these parameters in hand, we next estimated the *past* transcriptional state of each cell in our dataset (**Methods**), and sought to use these estimated states to link individual cells to one another across timepoints (Fig. 3a). Specifically, for each cell B (*e.g.* a cell from the 2 hr timepoint), we used a recently developed alignment method^33^ to identify a cell A profiled at an earlier timepoint (*e.g.* a cell from the 0 hr timepoint), wherein A’s current state was closest to B’s estimated past state. In this framework, A can be regarded as the parent state of B. Applying this strategy to each of the five intervals comprising our experiment, we constructed a set of linkages spanning the entire dataset and time course (Fig. 3b).

**Fig. 3.**
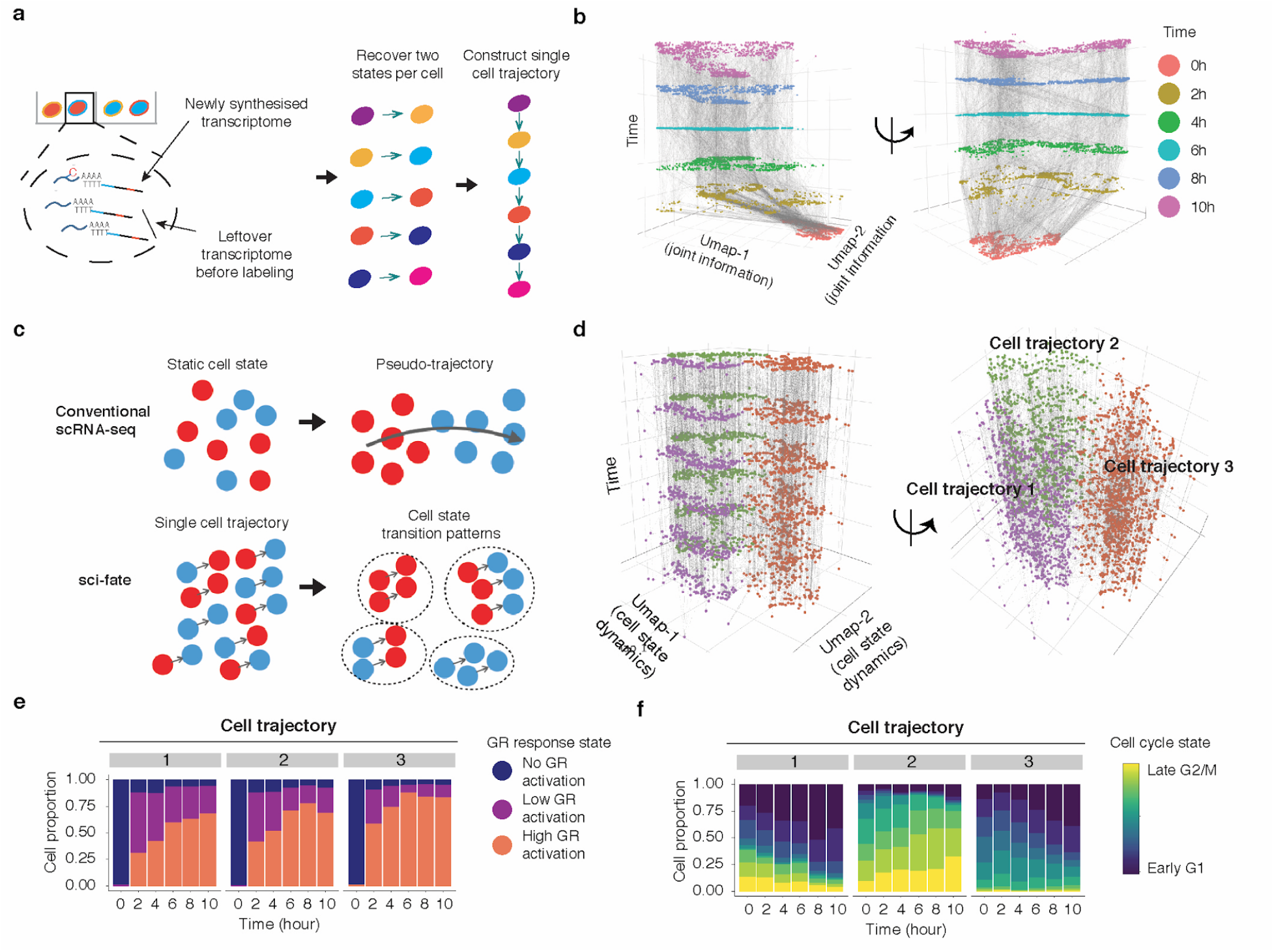
Inferring single cell transcriptional dynamics with sci-fate. (**a**) Schematic of approach for linking cells based on estimated past transcriptional states to reconstruct single cell transition trajectories. (**b**) 3D plot of all cells. The x and y coordinates correspond to the joint information UMAP space shown in the rightmost panel of Fig. 1e. The z coordinate as well as colors correspond to DEX treatment time. Linked parent and child cells are connected with grey lines. (**c**) Schematic comparing conventional scRNA-seq and sci-fate for cell trajectory analysis. (**d**) Similar to panel **b**, except the x and y coordinates correspond to the UMAP space based on the single cell transition trajectories across the six time points. (**e-f**) Barplots showing the contributions of the 3 GR response states (**e**) and the 9 different cell cycle states (**f**) to each of three cell trajectory clusters.

A key contrast with conventional pseudotime is that with sci-fate, each cell is now characterized not only by its present state, but also by specific linkages to a series of distinct cells matching its predicted past and/or future states (Fig. 3c). To evaluate whether there is structure to these mini-trajectories, we applied UMAP and unsupervised clustering, which resulted in three distinct trajectory clusters (Fig. 3d). To annotate these, we checked the proportions of each of the aforementioned three GR response states and nine cell cycle states in each of them, as a function of time. As expected, all three trajectories exhibited a rapid transition from no GR activation to low/high GR activation (Fig. 3e). However, each trajectory appears to correspond to a different starting point with respect to the cell cycle (Fig. 3f). Trajectory 1 corresponds to cells that transition from G2/M to G1 phase over the course of the 10 hr experiment. Trajectory 2 corresponds to cells that transition from late S phase to G2/M phase over the course of the experiment. Finally, trajectory 3 corresponds to cells that transition from G1 to either S phase or G1 arrest over the course of the experiment. The inference of G1 arrest subsequent to DEX treatment is consistent with the dynamics of cell state proportions in this experiment as well as with previous research^56,57^.

### Inferred single cell state transition links recapitulate expected dynamics

We next sought to evaluate whether the distribution of cell state transitions inferred by sci-fate are consistent with the expected dynamics. We assigned each cell into one of the 27 states (3 GR response x 9 cell cycle states) and computed a cell state transition network (Fig. 4a), with the assumption that the cell state transitions in this experiment follow a Markov process with a distribution that does not change over time. This assumption is validated in part by the observation that the distribution of predicted cell state transitions estimated from part of the data (0 hrs to 6 hrs) are highly correlated with similar estimates from the full data (Extended Data Fig. 6a). Although the cell state proportions were highly dynamic over the ten hour experiment (Extended Data Fig. 6b), the cell state transition network accurately predicted these proportions across all later timepoints based on the proportions from the first timepoint alone (Extended Data Fig. 6c). Consistent with the DEX treatment, transitions are highly biased from no/low to high GR activation states, as well as from G1 to S, S to G2/M, and G2/M to G1 phase of the cell cycle (Fig. 4a). As a control analysis, the cell state transition network based on randomly permuted cell state transition links failed to recapitulate these expected dynamics (Extended Data Fig. 6d).

**Fig. 4.**
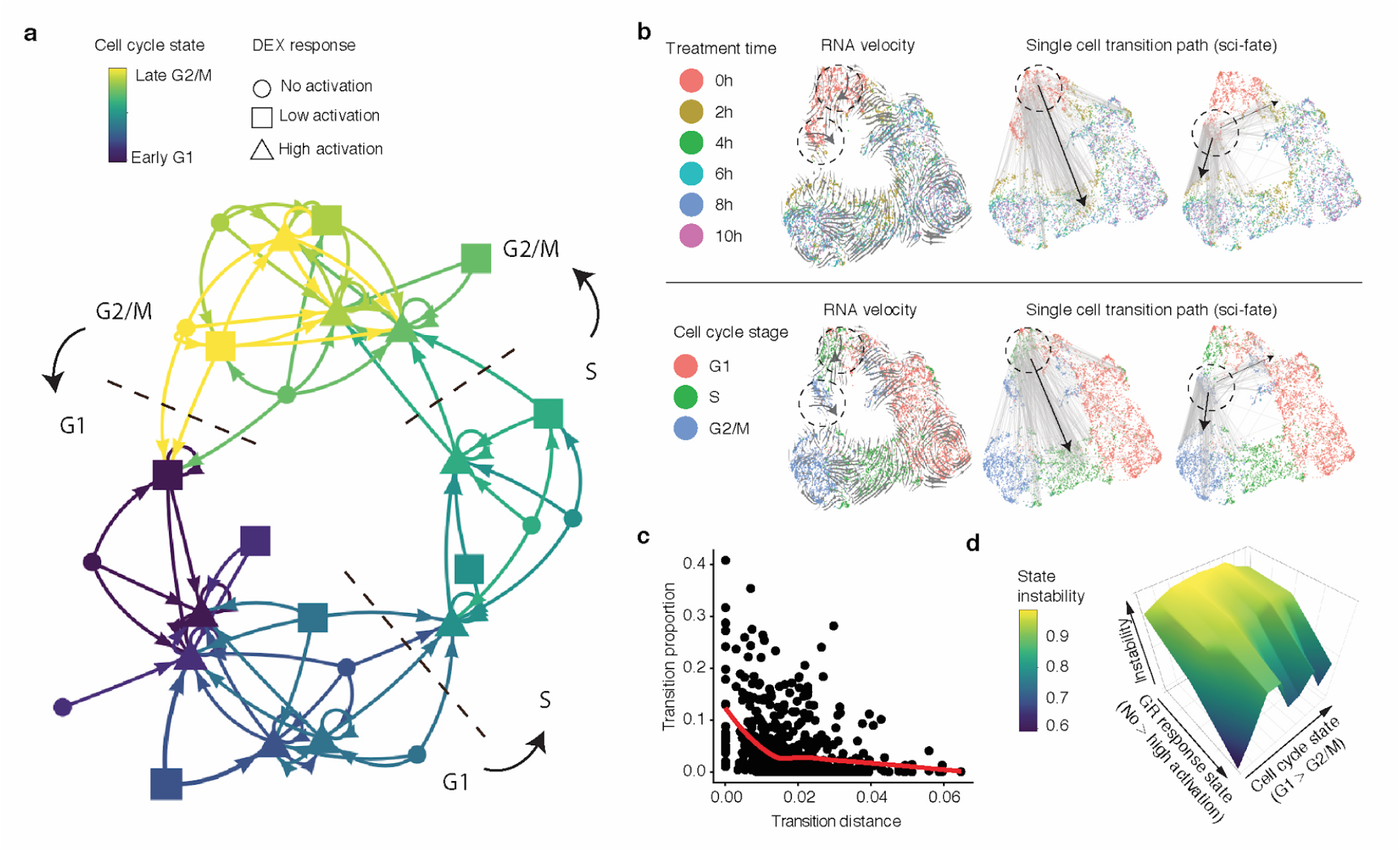
Constructing a state transition network for GR response and cell cycle. (**a**) Cell state transition network. The nodes are 27 cell states characterized by combinations of cell cycle and GR activation states. The links represents frequent cell state transition trajectories (transition proportion > 10%) between cell states. This threshold for defining a link corresponds to approximately two standard deviations from the mean transition proportion calculated after permuting cell transition links. (**b**) The x and y coordinates correspond to the joint information UMAP space shown in the rightmost panel of Fig. 1e, colored by DEX treatment time (top) or inferred cell cycle state (bottom). Grey lines represent inferred cell state transition directions by RNA velocity^58^ (left) or cell state transition links between parent and child cells in sci-fate (middle: cell state transition links starting from cells at S phase and no GR activation stage; right: cell state transition links starting from cells at G2/M phase and no GR activation stage). Black arrows show main cell state transition directions. (**c**) Scatter plot showing the relationship between transition distance (Pearson’s distance) and transition proportion, together with the red LOESS smoothed line by ggplot2^55^. (**d**) 3D plot showing the cell state stability landscape. X-axis represents GR response states (from no to low to high activation state). Y-axis represents the cell cycle states ordered from G1 to G2/M. Z-axis represents cell state instability, defined as the proportion of cells inferred to be moving out of a given state between time points.

The 27 states shown in Fig. 4a each correspond to subsets of cells whose transcriptomes are similar, making use of the joint information provided by distinguishing between old (> 2 hrs) vs. new (< 2 hrs) transcripts. Importantly, the distribution of transitions are inferred, rather than explicitly known, but supported by the fact that they correspond to expected phenomena, *e.g.* irreversible progression through GR response, as well as irreversible progression through the cell cycle. As an example, S phase cells without GR activation (0 hrs treatment) mostly transit into cell state in S phase with GR activation (2 hrs treatment), while G2/M phase cells with no GR activation (0 hrs treatment) mostly transit to G2/M or G1 phase cells with GR activation (2 hrs treatment) (Fig. 4b, upper). For comparison, reanalyzing the data with the RNA velocity method^58^, which infers transcriptional dynamics in single cell data from the proportion of intronic vs. exonic reads, failed to recover these expected patterns (Fig. 4b, lower). This could be because RNA velocity depends on the relative coordinates of cells in a low dimensional space, which is not a limitation of sci-fate.

Can we use this framework to better understand the characteristics of transcriptional states that govern their dynamics? As a first approach, we calculated the pairwise Pearson’s distance between the aggregated transcriptomes of each of the 27 states. As expected, the greater the distance between any pair of states, the lower the proportional representation of that transition in the network (Spearman’s correlation coefficient = −0.38; Fig. 4c). As a second approach, we computed “instability” as the proportion of cells inferred to be moving out of a given state between timepoints (Fig. 4d). As expected, states corresponding to no GR activation were the least stable by this metric. Furthermore, amongst high GR activation states, states corresponding to early G1 were the most stable. These representations of the data are consistent with the transition network, wherein the states corresponding to high GR activation and early G1 are a frequent “destination” of all nearby states (purple triangles in Fig. 4a).

## Discussion

Experimental methods that recover not only the current state of any given cell, but also its vector, are distinct and potentially more powerful than computational methods for inferring such vectors, *e.g.* pseudotime. To that end, we developed sci-fate, a novel method for concurrently profiling the whole and newly synthesized transcriptome in each of many single cells. In applying sci-fate to a model system of cortisol response, we found that the joint analysis of whole and newly synthesized single cell transcriptomes enabled greater discrimination of cell states than was possible with whole transcriptomes alone. Most notably, it became straightforward to distinguish between the dynamic transcriptional modules underlying GR activation vs. progression through the cell cycle. By analyzing covariance between TF expression and *new* RNA synthesis across many single cells, we identified regulatory links between 27 TFs and nearly one thousand target genes. These separated into several modules, including the GR response, cell cycle and others, reflecting cellular processes that were heterogeneous across this population of cells and that appeared to operate largely independently of one another. We were also able to infer the past state of each single cell in the experiment, and to use links between cells based on these inferences to construct a cell state transition network. Thus, sci-fate could in principle help overcome limitations of conventional single-cell RNA-seq when inferring causal regulatory networks and should help drive developmental of computational methods towards this end^59^.

Sci-fate captures information that is analogous to RNA velocity^58^, which distinguishes ‘older’ vs. ‘newer’ transcripts based on their splicing status. On one hand, RNA velocity is more straightforward than sci-fate, as it makes use of information that is indirectly captured by many single cell profiling technologies, whereas sci-fate requires S4U labeling steps that cannot necessarily be used in all contexts. On the other hand, sci-fate lends itself to experimental control in a way that RNA velocity does not, as the timing and length of S4U labeling can be specified whereas with RNA velocity it is a product of endogenous splicing dynamics. Furthermore, as we show, gene-specific mRNA degradation rates as well as the past transcriptional state of each cell can inferred, which may enable the quantitative analysis of cells with complex transcriptional histories and futures (*e.g.* multiple dynamics modules).

Because it is based on combinatorial indexing, sci-fate will be straightforward to scale to millions of cells^60^. It is also potentially compatible with concurrent profiling of the epigenomes from the same cells^61^, although ideally, we would be able to profile not only nascent transcription but also nascent epigenetic events. A major limitation of sci-fate is that S4U labeling experiments are generally performed *in vitro*. However, recent studies have shown that S4U can be used in conjunction with transgenic *UPRT*-expressing mice to stably label cell type-specific nascent RNA transcription *in vivo^62–64^*, suggesting that sci-fate, with further optimizations to enhance S4U incorporation and detection rate, can potentially be used to profile single transcriptional dynamics *in vivo* and at scale.

---

**Extended Data Fig. 1.**
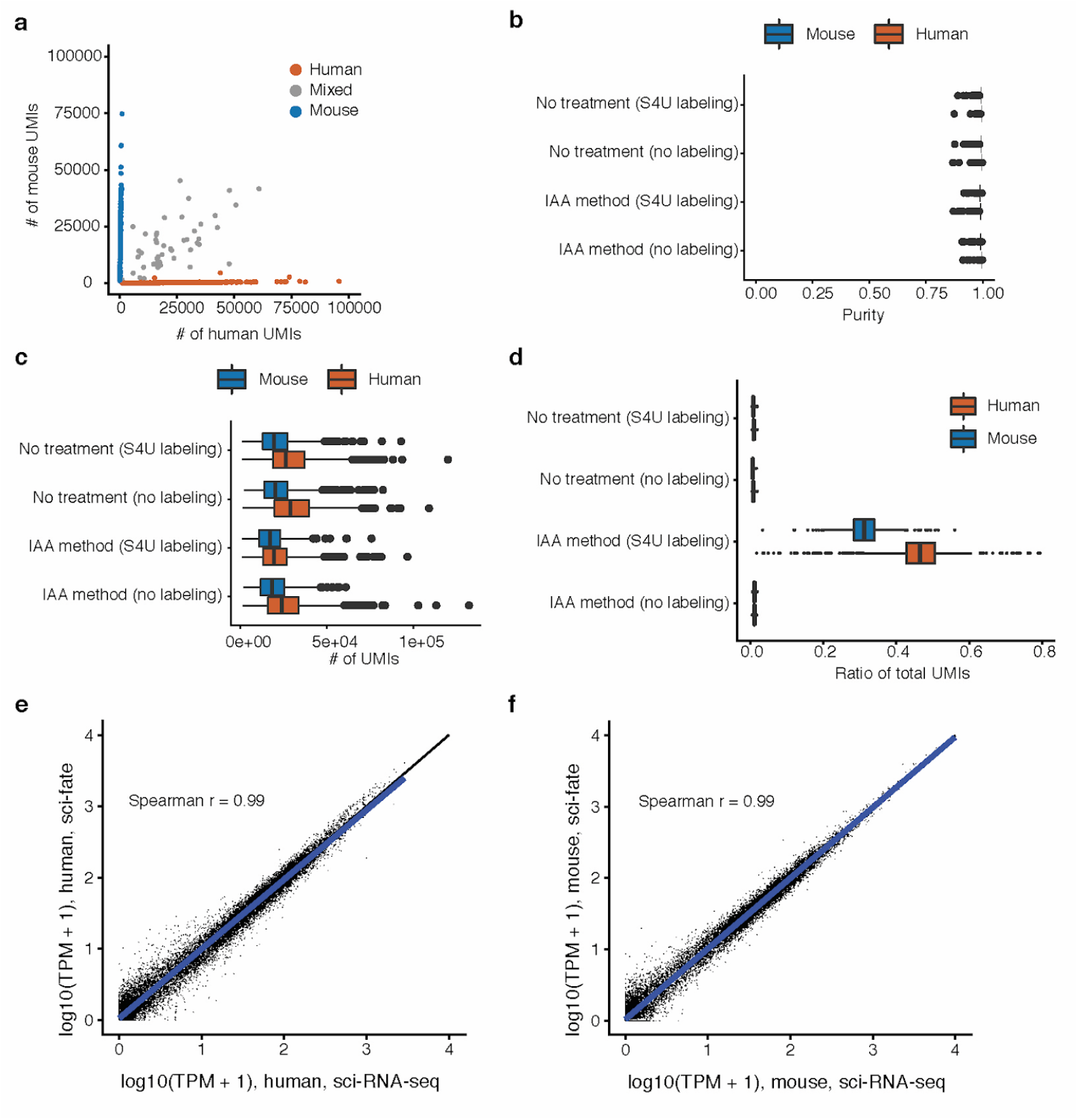
Performance and QC-related analyses for sci-fate. (**a**) Scatter plot of mouse (NIH/3T3) vs. human (HEK293T) UMI counts per cell in sci-fate. (**b-d**) Boxplot showing the proportion of reads mapping to the expected species (b), number of UMIs (c) and ratio of S4U labeled reads (d) per cell from HEK293T (n = 932) and NIH/3T3 (n = 438) cells. For all box plots: thick horizontal lines, medians; upper and lower box edges, first and third quartiles, respectively; whiskers, 1.5 times the interquartile range; circles, outliers. (**e-f**) Spearman’s correlation between gene expression measurements in aggregated profiles of HEK293T (e) and NIH/3T3 cells (f) from sci-fate (y-axis) vs. sci-RNA-seq (x-axis) cells.

**Extended Data Fig. 2.**
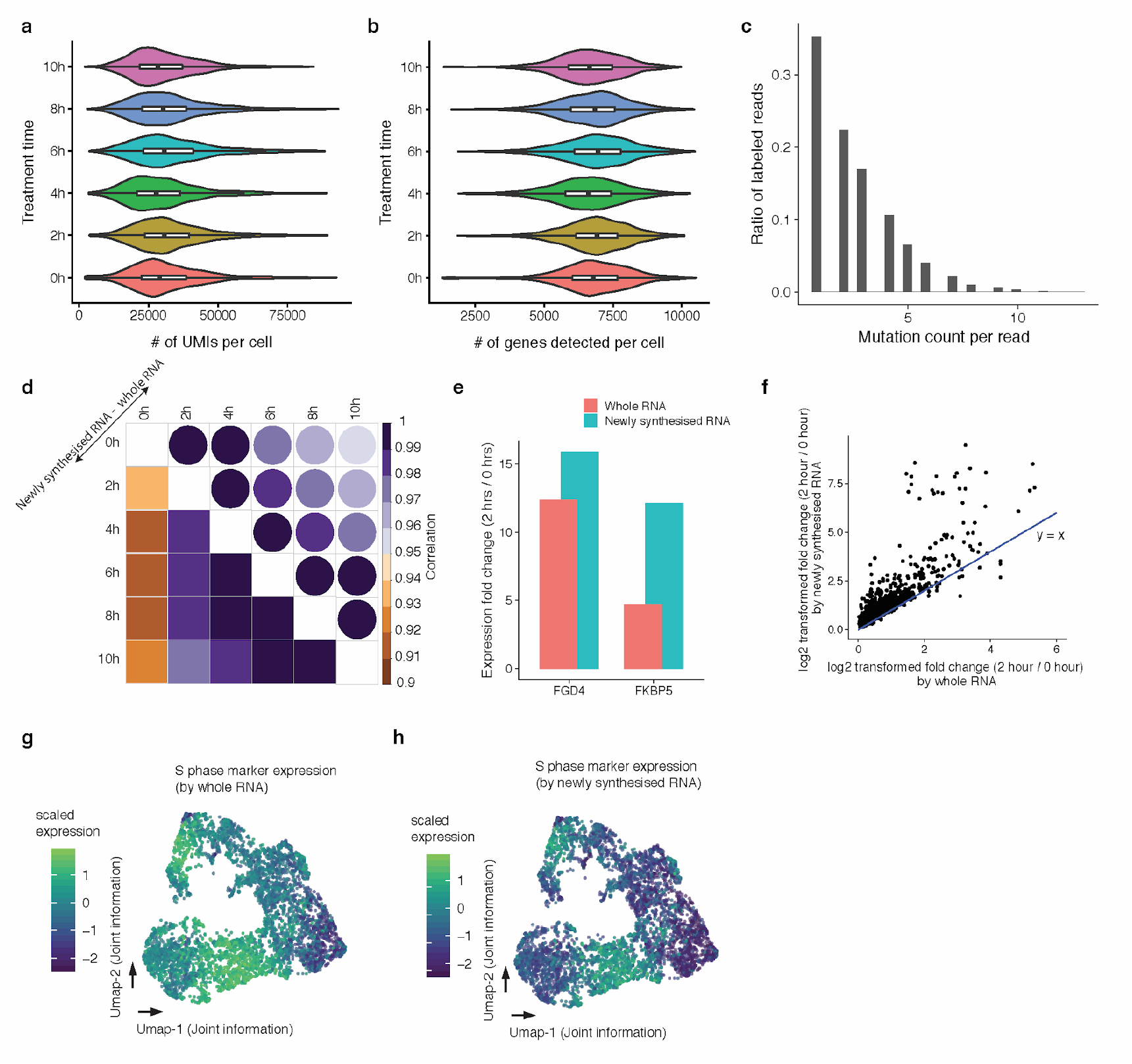
Performance of sci-fate on dexamethasone-treated A549 cells. (**a, b**) Violin plot showing the number of UMIs (a) and genes (b) per cell in each of six treatment conditions. For all box plots: thick horizontal lines, medians; upper and lower box edges, first and third quartiles, respectively; whiskers, 1.5 times the interquartile range; circles, outliers. (**c**) Histogram showing the distribution of of T > C mutation counts across labeled reads, estimated using reads from 100 randomly sampled cells and normalized by the total read number. (**d**) Correlation plot showing the Pearson’s correlation between different treatment conditions, using either whole transcriptomes (upper right; circles) or newly synthesized transcriptomes (bottom left; squares). (**e**) Bar plot showing the expression fold-change before and after 2 hour DEX treatment for *FGD4* and *FKBP5*, calculated using either whole RNA and newly synthesised RNA. (**f**) Scatter plot showing the expression fold-change before and after 2 hour DEX treatment for differentially expressed (DE) genes identified using either whole transcriptomes or newly synthesized transcriptomes. The blue line represent y = x line. (**g-h**) UMAP visualization of A549 cells via joint information of the whole and newly synthesized transcriptomes, colored by normalized expression of S phase marker genes in the whole (g) and newly synthesized (h) transcriptomes. UMI counts for these genes are scaled for library size, log-transformed, aggregated and then mapped to Z-scores.

**Extended Data Fig. 3.**
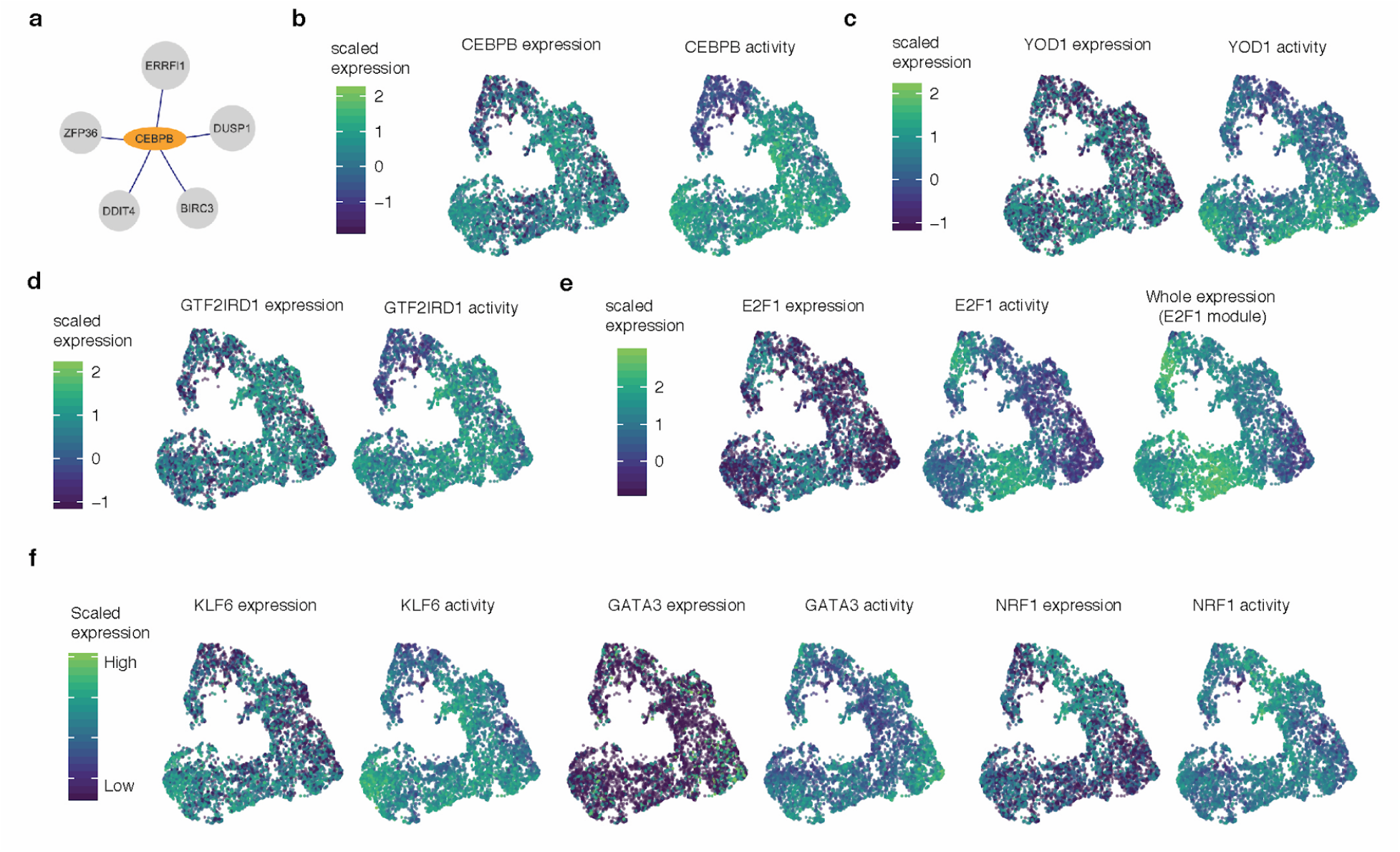
TF modules driving cell state transitions in DEX-treated A549 cells. (**a**) Identified gene targets (grey) of *CEBPB* (orange). Only links with regularized correlation coefficients from LASSO > 0.06 are shown. (**b**) UMAP visualization of A549 cells via joint information of the whole and newly synthesized transcriptomes, colored by *CEBPB* expression (left) and activity (right). (**c**) similar to (b), colored by the *YOD1* expression (left) or activity (right). (**d**) similar to (b), colored by the *GTF2IRD1* expression (left) or activity (right). (**e**) similar to (b), colored by the *E2F1* expression (left), activity (middle) or aggregated whole transcriptome expression of *E2F1* linked genes (right). (**f**) similar to (b), colored by the expression and activity of *KLF6* (left two panels), *GATA3* (middle two panels) or *NRF1* (right two panels).

**Extended Data Fig. 4.**
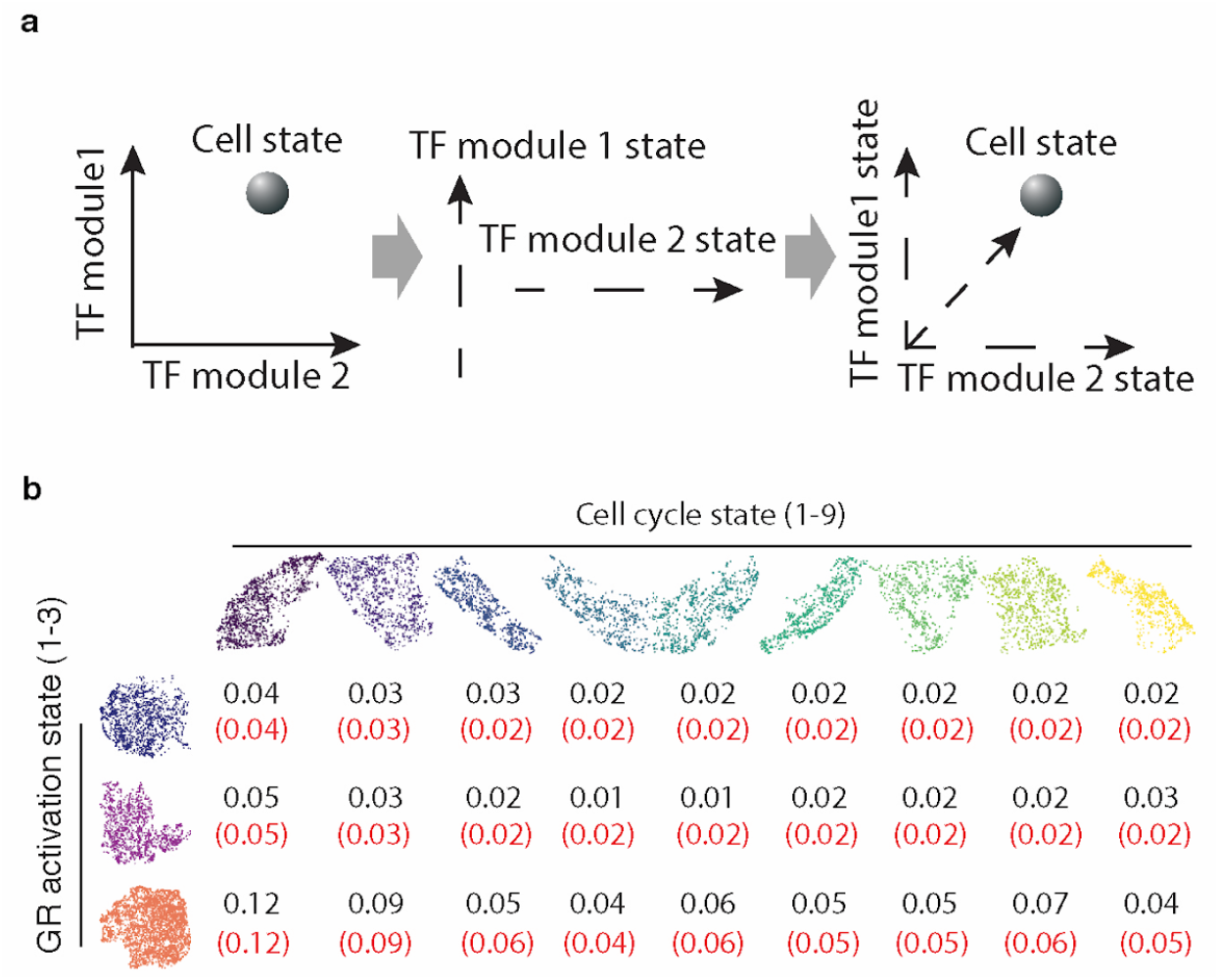
Twenty-seven cell states defined by combinations of TF module-defined states. (**a**) Schematic of strategy for characterizing cell states as the combination of TF modules. (**b**) Supplementary Table showing the observed proportion of cells (black numbers) falling into each of 27 cell states that correspond each correspond to one of 3 states defined by GR response (rows) and one of 9 states defined by the cell cycle module (columns). The red numbers in parentheses correspond to the expected proportions, assuming that the distributions are independent.

**Extended Data Fig. 5.**
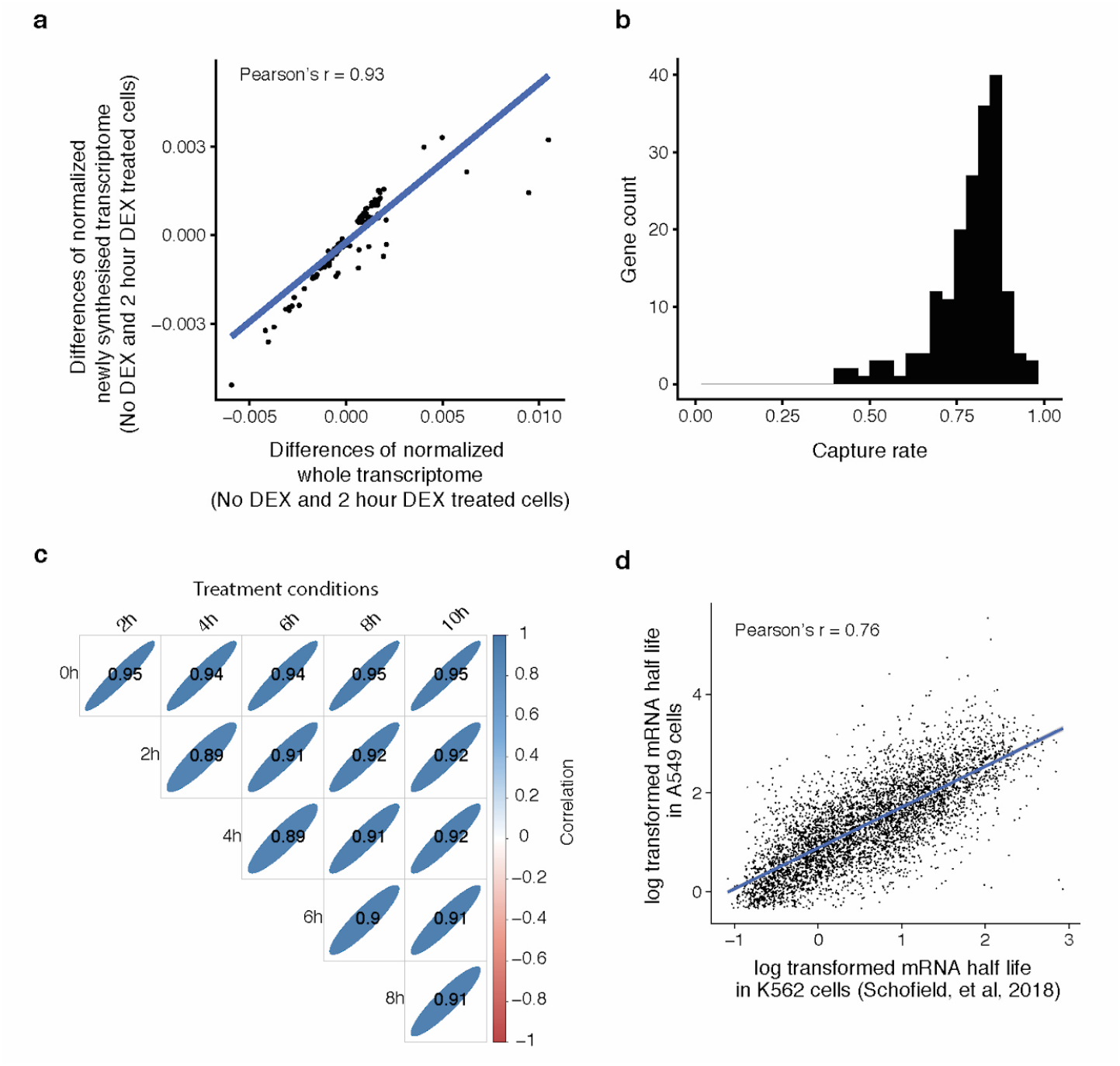
Estimating the rate of new RNA detection and the rates of RNA degradation. (**a**) Scatter plot showing differences between the normalized transcriptomes with no DEX treatment vs. 2 hours DEX treatment for each of 186 genes exhibiting the largest differences in new transcription between the two conditions. X-axis shows differences between the conditions in the whole transcriptome. Y-axis shows differences between the conditions in the newly synthesized transcriptome. Blue line is the linear regression line. Both whole transcriptome and newly synthesized transcriptome of each time point are normalized by the library size of whole transcriptome. (**b**) Histogram showing the distribution of the estimated capture rate of newly synthesised mRNA for each of 186 genes. **(c)** Correlation plot showing the Pearson’s correlation (r) of log-transformed gene degradation rate between treatment conditions. Positive correlations are displayed in blue and negative correlations in red. The shape of the ellipses are illustrative, determined by the correlation coefficients shown. (**d**) Scatter plot showing the correlation between estimated per-gene mRNA half lives (log transformed) in K562 cells from the literature (x-axis)^9^ vs. as estimated by sci-fate in A549 cells (y-axis).

**Extended Data Fig. 6.**
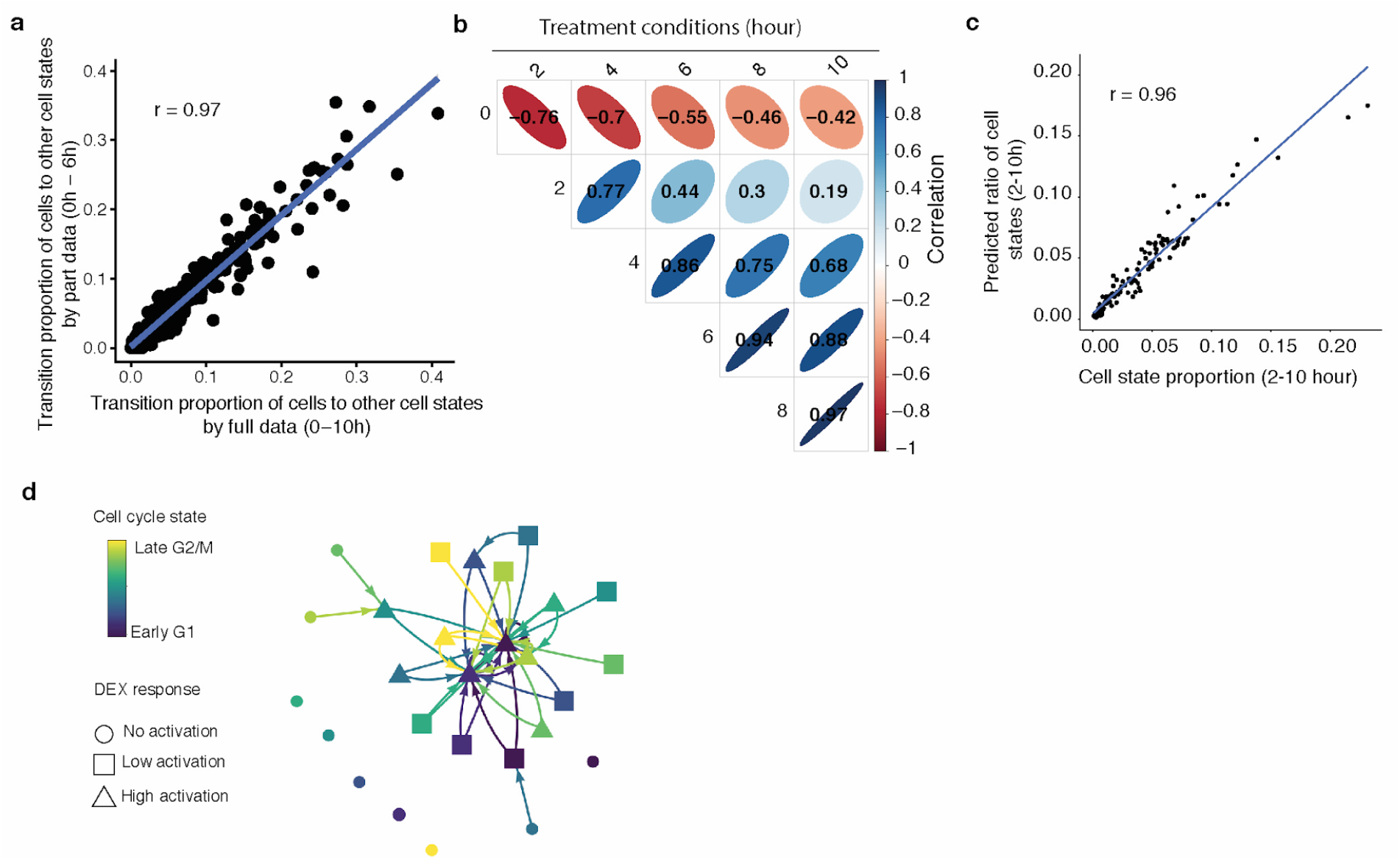
Constructing a state transition network for GR response and cell cycle. (**a**) Scatter plot showing the correlation of distribution of inferred cell state transitionscalculated by the full data (x-axis; 0-10 hours) vs. partial data (y-axis; 0-4 hours). The blue line represents the linear regression. (**b**) Correlation plot showing the Pearson’s correlation (r) of cell state proportions between treatment conditions. Positive correlations are displayed in blue and negative correlations in red. The shape of the ellipses are illustrative, determined by the correlation coefficients shown. (**c**) Scatter plots showing the observed vs. predicted cell state proportions for the 2, 4, 6, 8 and 10 hr DEX treatment groups. The predicted values are based on using a cell state transition network constructed from the full data. (**d**) Cell state transition network similar to Fig. 4a, but after permuting the parent and child cell links. During permutation, each child cell of each time point is linked to a randomly selected parent cell from the immediately preceding time point.

## Endnotes

### Acknowledgements

We thank members of the Shendure labs for helpful discussions, particularly Xingfan Huang, Ronnie Blecher, Beth Martin, Flo Chardon and Ruolan Qiu.

### Author contributions

J.S. and J.C. designed the research; J.C. developed technique and performed experiments with assistance from F.S.; J.C. performed computation analysis with suggestions from W.Z. and C.T.; J.S. and J.C. wrote the paper.

## Materials and Methods

### Mammalian cell culture

All mammalian cells were cultured at 37°C with 5% CO_2_, and were maintained in high glucose DMEM (Gibco cat. no. 11965) for HEK293T and NIH/3T3 cells or DMEM/F12 medium for A549 cells, both supplemented with 10% FBS and 1X Pen/Strep (Gibco cat. no. 15140122; 100U/ml penicillin, 100 µg/ml streptomycin). Cells were trypsinized with 0.25% typsin-EDTA (Gibco cat. no. 25200-056) and split 1:10 three times per week.

### Sample processing for sci-fate

A549 cells were treated with 100 nM DEX for 0 hrs, 2 hrs, 4 hrs, 6 hrs, 8 hrs or 10 hrs. Cells in all treatment conditions were incubated with 200uM S4U for the last two hours before cell harvest. For HEK293T and NIH/3T3 cells, cells were incubated with 200uM S4U for 6 hours before cell harvest.

All cell lines (A549, HEK293T and NIH/3T3 cells) were trypsinized, spun down at 300x**g** for 5 min (4°C) and washed once in 1X ice-cold PBS. All cells were fixed with 4ml ice cold 4% paraformaldehyde (EMS) for 15 min on ice. After fixation, cells were pelleted at 500x**g** for 3 min (4°C) and washed once with 1ml PBSR (1 x PBS, pH 7.4, 1% BSA, 1% SuperRnaseIn, 1% 10mM DTT). After wash, cells were resuspended in PBSR at 10 million cells per ml, and flash frozen and stored in liquid nitrogen. Paraformaldehyde fixed cells were thawed on 37°C water bath, spun down at 500x**g** for 5 min, and incubated with 500ul PBSR including 0.2% Triton X-100 for 3min on ice. Cells were pelleted and resuspended in 500ul nuclease free water including 1% SuperRnaseIn. 3ml 0.1N HCl were added into the cells for 5min incubation on ice ^25^. 3.5ml Tris-HCl (pH = 8.0) and 35ul 10% Triton X-100 were added into cells to neutralize HCl. Cells were pelleted and washed with 1ml PBSR. Cells were resuspended in 100ul PBSR. 100ul PBSR with fixed cells were incubated with mixture including 40ul Iodoacetamide (IAA, 100mM), 40ul sodium phosphate buffer (500mM, pH = 8.0), 200ul DMSO and 20ul H2O, at 50°C for 15min. The reaction was quenched by 8ul DTT (1M) and 8.5ml PBS^65^. Cells were pelleted and resuspended in 100ul PBSI (1 x PBS, pH 7.4, 1% BSA, 1% SuperRnaseIn). For all later washes, nuclei were pelleted by centrifugation at 500x**g** for 5 min (4°C).

The following steps are similar with sci-RNA-seq protocol with paraformaldehyde fixed nuclei ^19,20^. Briefly, cells were distributed into four 96-well plates. For each well, 5,000 nuclei (2 µL) were mixed with 1 µl of 25 µM anchored oligo-dT primer (5′-ACGACGCTCTTCCGATCTNNNNNNNN[10bp index]TTTTTTTTTTTTTTTTTTTTTTTTTTTTTTVN-3′, where “N” is any base and “V” is either “A”, “C” or “G”; IDT) and 0.25 µL 10 mM dNTP mix (Thermo), denatured at 55°C for 5 min and immediately placed on ice. 1.75 µL of first-strand reaction mix, containing 1 µL 5X Superscript IV First-Strand Buffer (Invitrogen), 0.25 µl 100 mM DTT (Invitrogen), 0.25 µl SuperScript IV reverse transcriptase (200 U/μl, Invitrogen), 0.25 μL RNaseOUT Recombinant Ribonuclease Inhibitor (Invitrogen), was then added to each well. Reverse transcription was carried out by incubating plates at the following temperature gradient: 4°C 2 minutes, 10°C 2 minutes, 20°C 2 minutes, 30°C 2 minutes, 40°C 2 minutes, 50°C 2 minutes and 55°C 10 minutes. All cells (or nuclei) were then pooled, stained with 4’,6-diamidino-2-phenylindole (DAPI, Invitrogen) at a final concentration of 3 μM, and sorted at 25 nuclei per well into 5 μL EB buffer. Cells were gated based on DAPI stain such that singlets were discriminated from doublets and sorted into each well. 0.66 μl mRNA Second Strand Synthesis buffer (NEB) and 0.34 μl mRNA Second Strand Synthesis enzyme (NEB) were then added to each well, and second strand synthesis was carried out at 16°C for 180 min. Each well was then mixed with 5 μL Nextera™ TD buffer (Illumina) and 1 μL i7 only TDE1 enzyme (25 nM, Illumina, diluted in Nextera™ TD buffer), and then incubated at 55°C for 5 min to carry out tagmentation. The reaction was stopped by adding 12 μL DNA binding buffer (Zymo) and incubating at room temperature for 5 min. Each well was then purified using 36 uL AMPure XP beads (Beckman Coulter), eluted in 16 μL of buffer EB (Qiagen), then transferred to a fresh multi-well plate.

For PCR reactions, each well was mixed with 2μL of 10 μM P5 primer (5′-AATGATACGGCGACCACCGAGATCTACAC[i5]ACACTCTTTCCCTACACGACGCTCTTCCGATCT-3′; IDT), 2 μL of 10 μM P7 primer (5′-CAAGCAGAAGACGGCATACGAGAT[i7]GTCTCGTGGGCTCGG-3′; IDT), and 20 μL NEBNext High-Fidelity 2X PCR Master Mix (NEB). Amplification was carried out using the following program: 72°C for 5 min, 98°C for 30 sec, 18-22 cycles of (98°C for 10 sec, 66°C for 30 sec, 72°C for 1 min) and a final 72°C for 5 min. After PCR, samples were pooled and purified using 0.8 volumes of AMPure XP beads. Library concentrations were determined by Qubit (Invitrogen) and the libraries were visualized by electrophoresis on a 6% TBE-PAGE gel. Libraries were sequenced on the NextSeq™ 500 platform (Illumina) using a V2 150 cycle kit (Read 1: 18 cycles, Read 2: 130 cycles, Index 1: 10 cycles, Index 2: 10 cycles).

### Read alignment and downstream processing

Read alignment and gene count matrix generation for the single cell RNA-seq was performed using the pipeline that we developed for sci-RNA-seq^10^ with minor modifications. Reads were first mapped to a reference genome with STAR/v2.5.2b^66^, with gene annotations from GENCODE V19 for human, and GENCODE VM11 for mouse. For experiments with HEK293T and NIH/3T3 cells, we used an index combining chromosomes from both human (hg19) and mouse (mm10). For the A549 experiment, we used human genome build hg19.

The single cell sam files were first converted into alignment tsv file using sam2tsv function in jvarkit^67^. Next, for each single cell alignment file, mutations matching the background SNPs were filtered out. For background SNP reference of A549 cells, we downloaded the paired-end bulk RNA-seq data for A549 cells from ENCODE^35^ (sampled name: ENCFF542FVG, ENCFF538ZTA, ENCFF214JEZ, ENCFF629LOL, ENCFF149CJD, ENCFF006WNO, ENCFF828WTU, ENCFF380VGD). Each paired-end fastq files were first adaptor-clipped using trim_galore/0.4.1^68^ with default settings, aligned to human hg19 genome build with STAR/v2.5.2b^66^. Unmapped and multiple mapped reads were removed by samtools/v1.3^69^. Duplicated reads were filtered out by MarkDuplicates function in picard/1.105 ^70^. De-duplicated reads from all samples were combined and sorted with samtools/v1.3^69^. Background SNPs were called by mpileup function in samtools/v1.3^69^ and mpileup2snp function in VarScan/2.3.9 ^71^. For HEK293T and NIH/3T3 test experiment, background SNP reference was generated in a similar pipeline above, with the aggregated single cell sam data from control condition (no S4U labeling and no IAA treatment condition).

For each single cell alignment file, all mutations with quality score <= 13 were removed. Mutations at the both ends of each reads were mostly due to sequencing errors, and thus also were filtered out. For each read, we checked if there are T > C mutations for sense strand or A > G mutations for antisense strand, and labeled these mutated reads as newly synthesized.

Each cell was characterized by two digital gene expression matrixes from the full sequencing data and newly synthesized RNA data as described above. Genes with expression in equal or less than 5 cells were filtered out. Cells with fewer than 2,000 UMIs or more than 80,000 UMIs were discarded. Cells with doublet score > 0.2 by doublet analysis pipeline Scrublet/0.2^72^ were removed.

The dimensionality of the data was first reduced with PCA (after selecting the top 2,000 genes with highest variance) on digital gene expression matrixes on either full gene expression data or the newly synthesized gene expression data by Monocle 3^2,73^. The top 10 PCs were selected for dimensionality reduction analysis with uniform manifold approximation and projection (UMAP/0.3.2), a recently proposed algorithm based on Riemannian geometry and algebraic topology to perform dimension reduction and data visualization^31^. For joint analysis, we combined top 10 PCs calculated on the whole transcriptome and top 10 PCs on the newly synthesized transcriptome for each single cell before dimension reduction with UMAP. Cell clusters were done via densityPeak algorithm implemented in Monocle 3^2,73^. We first performed UMAP analysis on joint information of all processed cells, and identified an outlier cluster (724 out of 7,404 cells). These cells were marked by high level expression of *GATA3*, a marker of differentiated cells^43^, and were filtered out before downstream analysis.

### Linking transcription factors (TFs) to their regulated genes

We sought to identify links between TFs and their regulated genes based on expression covariance. Cells with more than 10,000 UMIs detected, and genes with newly synthesis reads detected in more than 10% of all cells were selected. The full gene expression and newly synthesized gene count per cell were normalized by cell-specific library size factors computed on the full gene expression matrix by estimateSizeFactors in Monocle 3^2,73^, log transformed, centered, then scaled by scale() function in R. For each gene detected, a LASSO regression model was constructed with package glmnet^74^ to predict the normalized expression levels, based on the normalized expression of 853 TFs annotated in the “motifAnnotations_hgnc” data from package RcisTarget^36^, by fitting the following model:

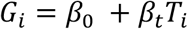

where *G*_*i*_ is the adjusted gene expression value for gene i. It is calculated by the newly synthesized mRNA count for each cell, normalized by cell specific size factor (*SG*_*i*_) estimate by estimateSizeFactors in Monocle 3^2,73^ on the full expression matrix of each cell, and log transformed:

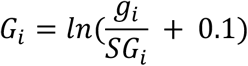

To simplify downstream comparison between genes, we standardize the response *G*_*i*_ prior to fitting the model for each gene *i* with the scale() function in R.

Similar with *G*_*i*_, *T*_*i*_is the adjusted TF expression value for each cell. It is calculated by the full TF expression count for each cell, normalized by cell specific size factor (*SG*_*i*_) estimate by estimateSizeFactors in Monocle 3^2,73^ on the full expression matrix of each cell, and log transformed:

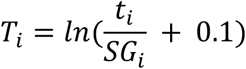

Prior to fitting, *T*_*i*_is standardized with the scale() function in R.

Although negative correlations between a TF’s expression and a gene’s new synthesis rate could reflect the activity of a transcriptional repressor, we felt that the more likely explanation for negative links reported by glmnet was mutually exclusive patterns of cell-state specific expression and TF activity. Thus during prediction, we excluded TFs with negatively correlated expression with a potential target gene’s synthesis rate, and also low correlation coefficient (<= 0.03) links. We identified a total of 6,103 links between TFs and regulated genes.

Our approach aims to identify TFs that may regulate each gene, by finding the subset that can be used to predict its expression in a regression model. However, a TF with expression correlated with a gene’s expression does not definitively mean that it is directly regulating that gene. To identify putatively direct targets within this set, we intersected the links with TFs profiled in ENCODE ChIP-seq experiments^35^. Out of 1,086 links with TFs characterized in ENCODE, 807 were validated by TF binding sites near gene promoters^34^, a 4.3-fold enrichment relative to background expectation (odds ratio for validation = 2.89 for links identified in LASSO regression vs. 0.67 for background, p-value < 2.2e-16, Fisher’s Exact test). Only gene sets with significant enrichment of the correct TF ChIP-seq binding sites were retained (Fisher’s Exact test, FDR of 5%), and further pruned to remove indirect target genes without TF binding data support. Ultimately, 591 links were retained by this approach.

To expand the set of validated TF-gene links, we further applied package SCENIC^36^, a pipeline to construct gene regulatory networks based on the enrichment of target TF motifs in the 10 kb window around genes’ promoters. Each co-expression module identified by LASSO regression was analyzed using cis-regulatory motif analysis using RcisTarget^36^. Only modules with significant motif enrichment of the correct TF regulator were retained, and pruned to remove indirect target genes without motif support. We filtered the TF-gene links by three correlation coefficient thresholds (0.3, 0.4 and 0.5), and combined all links validated by RcisTarget^36^. In total, there were 509 links validated this motif-based approach.

Combining both approaches, we identified a total of 986 TF-gene regulatory links by the covariance between TF expression and gene synthesis rate, validated by DNA binding data or motif analysis. To evaluate the possibility that the links were artifacts of regularized regression, we permuted the sample IDs of the TF expression matrix and performed the same analyses.No links were identified after this permutation.

### Ordering cells based on the activity of functional TF modules

To calculate TF “activity” in each cell, newly synthesized UMI counts for genes linked to each of the 27 TFs were scaled by library size, log-transformed, aggregated and then mapped to Z-scores. As TFs with highly correlated or anti-correlated activity suggest they may function in linked biological processes, we calculated the absolute Pearson’s correlation coefficient between each pair of TF activity, and based on this we clustered TFs by ward.d2 clustering method in package pheatmap/1.0.12^75^. Five functional TF modules were identified and annotated based on their functions.

To characterize the dynamics of cells in relation to potentially independent cellular processes, cells were ordered by the activity of cell cycle related TFs (TF module 1) or GR activity related TFs (TF module 3) with UMAP (metric = “cosine”, n_neighbors = 30, min_dist = 0.01). The cell cycle progression trajectory were validated by cell cycle gene markers in Seurat/2.3.4^33^. Three cell cycle phases were identified by densityPeak algorithm implemented in Monocle 3^2,73^, on the UMAP coordinates ordered by cell cycle TF modules. As each main cell cycle phase still showed variable TF activity and cell cycle marker expression, we segmented each phase to early/middle/late states by k-means clustering (k = 3), and recovered a total of nine cell cycle states. Three GR reponse states were identified by densityPeak algorithm implemented in Monocle 3^2,73^.

### Past transcriptome state recovery from sci-fate

To infer the past transcriptome state (*i.e.* the cell state before S4U labeling commenced), we assume mRNA half lives are consistent across different DEX treatment conditions. This assumption is further validated by self-consistency check later. Under this assumption, the partly degraded bulk transcriptome before the 2 hour S4U labeling should be the same between no DEX and 2 hour DEX treated cells. Thus, for any given gene, differences in whole transcriptomes (bulk) between these timepoints should be equal to differences in the newly synthesized transcriptomes (bulk), corrected by technique’s detection rate:

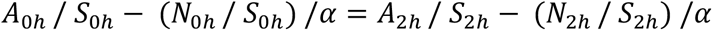

*A*_0*h*_ is the aggregated UMI count for all cells in no DEX treatment group; *S*_0*h*_ is the library size (total UMI count of cells) at no DEX treatment; *N*_0*h*_ is the aggregated newly synthesized UMI count for all cells in no DEX treatment group; *A*_2*h*_ is the aggregated UMI count for all cells in 2 hour DEX treatment group; *S*_2*h*_ is the library size (total UMI count of cells) in 2 hour DEX treatment group; *N*_2*h*_ is the aggregated newly synthesized UMI count for all cells in 2 hour DEX treatment group; α is the detection rate for sci-fate. In theory, one detection rate can be calculated for each gene. However, for genes with minor differences of newly synthesis rate between two conditions, the estimated α is dominated by noise. We thus selected genes showing higher differences in normalized newly synthesis rate between two conditions: we first tested a series of threshold for gene filtering and calculated the α for each gene. We then plotted the relationship between threshold and the ratio of genes with out-range α values (< 0 or > 1). We selected the threshold that was at the knee point of the plot, resulting in 186 genes selected. The differences in newly synthesized mRNA of these genes highly correlates with the differences in mRNA expression level (Pearson’s r = 0.93, Extended Data Fig. 5a), suggesting the new RNA detection rate is stable across genes (Extended Data Fig. 5b). We therefore moved forward with the median estimate (82%) of the proportion of newly synthesized RNA captured by sci-fate.

We next computed the mRNA degradation rate across each 2 hour interval. As the A549 cell population can be regarded stable without external perturbation, for 2 hour DEX treated cells, its past state (before 2 hour S4U labeling) should be the same as the 0 hour DEX treated cells. Expanding on this logic, the past state (before S4U labeling) for T = 0/2/4/6/8/10 hour DEX treated cells should be similar to the profiled T = 0/0/2/4/6/8 hour cells, respectively:

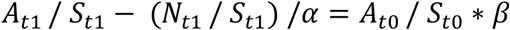

*A*_*t*1_ is the aggregated UMI count for all cells in t1; *S*_*t*1_ is is the library size (the total UMI count of cells) at t1; *N*_*t*1_ is the aggregated newly synthesized UMI count for all cells at t1; α is the estimated detection rate of sci-fate; *A*_*t*0_ is the aggregated UMI count for all cells in t0; *S*_*t*0_ is is the library size (the total UMI count of cells) at t0; *β* is 1 - gene specific degradation rate between t0 and t1, and is related with the mRNA half life *γ* by:

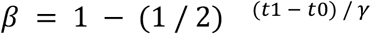

The gene degradation rate *β* can be calculated for each 2 hour interval of DEX treatment. As a self-consistency check mentioned above, the gene degradation rates are highly correlated across different DEX treatment times (Extended Data Fig. 5c). We therefore used the average degradation rate for each gene for downstream analysis.

With the overall sci-fate detection rate as well as per-gene degradation rates estimated, the past transcriptome state of each cell can be estimated by:

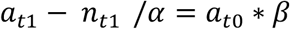

*a*_*t*1_ is the single cell UMI count in t1; *n*_*t*1_is the single cell newly synthesized UMI count at t1; α is the estimated detection rate of sci-fate; *β* is 1-gene specific degradation rate between t0 and t1. *a*_*t*0_ is the estimated single cell transcriptome in a past time point t0, with all negative values converted to 0.

### Linkage analysis to build single cell state trajectory

The goal of what we call here “linkage analysis” is to associate each cell with parent and child cells at different timepoints, *i.e.* single cell state trajectories. Our approach is based on a fact that the past transcriptome states (before 2 hour S4U labeling) of cells at t1 should share the same cell population distribution with the profiled transcriptome states of cells at t0 (2 hours earlier than t1), assuming there is no cell apoptosis. We thus applied a published manifold alignment strategy to identify common cell states between two data sets, based on common sources of variation^33^. As a result, whole transcriptomes from t0 cells and recovered past transcriptomes from t1 cells are aligned in the same UMAP space. This analysis is based on an assumption that for intermediate timepoints, we are oversampling the space of physiologically distinct states in this timecourse. Violation of this and other assumptions can be detected by outliers during alignment of the two data sets. For each cell A from t1, we selected its nearest neighbour in t0 as its parent state in the alignment UMAP space. Similarly, for each cell from t0, we selected its nearest neighbour in t1 as its child cell state. Of note, the link is not necessary to be bi-directional: the parent state of one cell may be linked to a different child cell. After the parent and child states were identified for each cell (except cells at the start and end time points), we then extend each cell trajectory by searching for the linked parent cell of each cell’s parent, and similarly the linked child cell of each cell’s child. Thus each single cell can be characterized by a single cell state transition path across all six time points spanning 10 hours. As multiple cells (>50) are profiled for each of the 27 defined cell state, stochastic cell state transition processes can also potentially be captured.

### Dimensionality reduction and clustering analysis

For dimensionality reduction on single cell transcriptomes, the top 5 PCs for full transcriptomes and top 5 PCs for newly synthesized transcriptomes were selected for each state, and combined in temporal order along single cell state trajectory for UMAP analysis. Main cell trajectory types were identified by density peak clustering algorithm^76^.

With cell state proportion at the beginning time point (0 hour treatment) and cell state transition probabilities estimated from the data, we first predicted the cell state distribution after 2 hours, assuming the cell state transitions in DEX treatment are cell-autonomous, time-independent, Markovian processes. Similarly, the cell state distribution at later time points can be predicted from the cell state distribution 2 hours before.

For RNA velocity analysis of these same data, single cell spliced/unspliced expression matrices were generated using the command line interface of velocyto/v.0.17^58^ with the default run_smartseq2 mode on single cell bam files. Cell transition direction inference were performed with an optimized scalable RNA velocity analysis toolkit scVelo/v.0.1.17 and scanpy/v.1.4.1 with default settings^58,77^.

### Cell state instability and cell state distance calculations

We defined cell state instability as the proportion of cells in a given state ‘moving’ to any other state at the next time point. To calculate cell state distances, we first sampled equal number (n = 50) of cells from each state, and separately aggregated the full transcriptome and newly synthesized transcriptome of sampled cells of that state (*i.e.* in this ‘joint transcriptome’, each gene is represented by two columns, one for the whole transcriptome and one for the newly synthesized transcriptome). The cell state distance is calculated as the Pearson’s correlation coefficient between the joint transcriptomes of two different states.

## Code Availability

Scripts for processing sci-fate sequencing were written in python and R with code available at https://github.com/JunyueC/scifate.

## Legends for Supplementary Table 1-3 (tables are provided separately as an Excel file)

Supplementary Table 1: Differentially expressed (DE) genes between 0 hour and 2 hour DEX timepoints.

Supplementary Table 2: Links between 27 TFs and 986 putatively regulated genes.

Supplementary Table 3: Gene degradation rate in each 2 hour time window estimated by sci-fate.

